# Data Aggregation at the Level of Molecular Pathways Improves Stability of Experimental Transcriptomic and Proteomic Data

**DOI:** 10.1101/076620

**Authors:** Nicolas Borisov, Maria Suntsova, Andrew Garazha, Ksenia Lezhnina, Olga Kovalchuk, Alexander Aliper, Elena Ilnitskaya, Maxim Sorokin, Mihkail Korzinkin, Vyacheslav Saenko, Yury Saenko, Dmitry G. Sokov, Nurshat M. Gaifullin, Kirill Kashintsev, Valery Shirokorad, Irina Shabalina, Alex Zhavoronkov, Bhubaneswar Mishra, Charles R. Cantor, Anton Buzdin

## Abstract

High throughput technologies opened a new era in biomedicine by enabling massive analysis of gene expression at both RNA and protein levels. Unfortunately, expression data obtained in different experiments are often poorly compatible, even for the same biological samples. Here, using experimental and bioinformatic investigation of major experimental platforms, we show that aggregation of gene expression data at the level of molecular pathways helps to diminish cross- and intra-platform bias otherwise clearly seen at the level of individual genes. We created a mathematical model of cumulative suppression of data variation that predicts the ideal parameters and the optimal size of a molecular pathway. We compared the abilities to aggregate experimental molecular data for the five alternative methods, also evaluated by their capacity to retain meaningful features of biological samples. The bioinformatic method OncoFinder showed optimal performance in both tests and should be very useful for future cross-platform data analyses.

## INTRODUCTION

Next generation sequencing (NGS), Microarray hybridization (MH) and high throughput proteomic techniques opened a new era in biomedicine by enabling large scale analysis of gene expression at both the RNA and protein levels [Kumar, 2016]. Multiple experimental platforms based on different principles and utilizing different reagents were developed for these tasks [Kumar, 2016]. According to the International Aging Research Portfolio, over eight billion dollars in government funding have been spent on research projects involving high throughput gene expression analysis since 1993 [Zhavoronkov, 2011]. This resulted in tens of thousands of publications. Unfortunately, gene expression data obtained using different experimental platforms are poorly compatible with each other even when obtained using the same biosamples. For example, a generally weak correlation between NGS and microarray gene expression data has been reported [Buzdin, 2014]. Therefore, a new data processing method is badly needed to enable data harmonization among different platforms and experiments [MAQC Consortium, 2006; Zhang, 2013].

Recently we showed that aggregation of gene expression data into molecular pathways, each containing dozens or hundreds of gene products, may help to solve the problem of poor data compatibility among different experimental platforms [Buzdin 2014a]. NGS and microarray data obtained for the same transcripts showed generally low correlation (<0.2) when examined at the level of individual genes. However, these correlations improved dramatically, up to 0.9, when activation of 90 molecular pathways was analyzed instead [Buzdin, 2014a]. The output measure was a Pathway Activation Strength (*PAS*), which positively reflects the degree of pathway activation. The *PAS* makes it possible to interrogate, quantitatively, processes such as molecular signaling, metabolism, DNA repair and cytoskeleton reorganization, based on gene expression data. These processes determine cell fate by governing growth, differentiation, proliferation, migration, survival and death [Diderich, 2016; Zhavoronkov, 2014]. Molecular modeling of intracellular pathways has been carried out for more than two decades [Kholodenko, 1999; Hanahan, 2000]. A plethora of molecular pathways have been discovered and catalogued, each containing different numbers of gene products [Haw, 2012; Nakaya et al., 2013]. Pathway activation strength was also found to be a better marker of human tissue types [Borisov, 2014; Lehznina, 2014] and tumor response to chemotherapy treatment [Zhu 2015;Venkova, 2015; Artemov, 2015]. Several approaches were published by us and others to assess the activation of signaling pathways, basing on large scale molecular data [Khatri, 2012; Buzdin, 2014b, Zhavoronkov, 2014]. These methods take into account different factors like the extent of differential gene expression, architecture of molecular pathways, and the roles of individual gene products in a pathway (e.g., activator/repressor) [Khatri, 2012; Buzdin, 2014b]. For example, a method we used to minimize discrepancies between the NGS and microarray platforms, termed OncoFinder, relies on differential gene expression and the known roles in a pathway, but does not take into account pathway architecture, i.e. the position of a gene product in a pathway [Buzdin, 2014b].

In spite of this progress it is not known why data aggregation improves expression information stability and what factors influence it. It is also unclear which bioinformatic algorithms provide better *PAS* outputs for cross-platform data stability. Additionally, *PAS* algorithms have not yet been applied to the high throughput proteomic data.

In this study, we applied data aggregation methods to transcriptomic information obtained using the Affymetrix HG U133 Plus 2.0, the Illumina HT12 bead array, the Agilent 1M array, the llumina Genome Analyzer platforms, and to proteomic data from the Orbitrap Velos and XL mass spectrometer platforms. We confirmed that for both transcriptomic and proteomic expression levels, the *PAS* approach provided more stable results than the expression of individual genes. To explain this phenomenon, we created a biomathematical model simulating error acquisition in individual gene expression and in *PAS*-based approaches. In agreement with the experimental data, in the mathematical model *PAS* methods produced significantly more stable results under a majority of conditions. This model also predicts the optimal size of a molecular pathway and ideal parameters of the normalizing (control) set of gene expression data.

To test the predictions further of the biomathematical model, we designed a new experimental gene expression array using the CustomArray microchip platform (USA) enabling direct electrochemical synthesis of oligonucleotide probes on a blank array. We compared results for the seven human kidney cancer tissue samples independently profiled by the two laboratories on the this customized array and on the commercial Illumina HT12 bead array platform. In agreement with the theoretical model, gene expression features differed significantly among the platforms for the same biosamples, while *PAS* values remained highly correlated. Therefore, gene expression data aggregated at the *PAS* level appears to be the method of choice for cross-platform data comparisons, including both transcriptomic and proteomic approaches.

We next explored the capacity of five most popular *PAS* calculation methods, OncoFinder [Buzdin, 2014b], TAPPA (Topology analysis of pathway phenotype association) [Gao, 2007], Topology-Based Score (TBScore) [Ibrahim, 2012], Pathway-Express [Draghici, 2007], and SPIA (Signal pathway impact analysis) [Tarca, 2009] to generate stable and biologically relevant data. We used the MicroArray Quality Control (MAQC) dataset [MAQC Consortium, 2006] including expression data for four biological samples profiled in fifteen replicates on major commercial microarray platforms. The abilities of the various *PAS* methods to increase correlation between transcriptomic features of the same biosamples examined using different experimental platforms were tested. We also checked whether the *PAS* methods were able to retain biological features after data harmonization using a generally accepted cross-platform harmonization procedure XPN [Shabalin, 2008]. We found that the OncoFinder method showed the optimal performance in both tests.

## RESULTS

### Cross-platform processing of transcriptomic and proteomic data

We processed transcriptomic and proteomic data to establish pathway activation strength (PAS) profiles corresponding to intracellular molecular pathways. The analysis included 271 molecular pathways (Supplementary dataset S1). For PAS measurements, we applied the OncoFinder method which was previously shown to diminish the cross-platform variation between the MH and NGS data [Buzdin, 2014a]. OncoFinder has previously been applied to many human and non-human systems including cell culture, leukemia and solid cancers, fibrosis, asthma, Hutchinson Gilford and Age-Related Macular Degeneration Disease [Makarev, 2016; Artcibasova, 2016; Alexandorva, 2016; Lebedev, 2015]. The PAS for a given pathway (*p)* is calculated as: 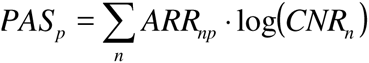 [Buzdin, 2014b], where the functional role of the n^th^ gene product in the pathway is indicated by the *activator/repressor role* (*ARR*), which equals 1 for an activator, – 1 for a repressor, and intermediate values −0,5; 0,5 and 0 for gene products having intermediate repressor, activator, or unknown roles, respectively. The *CNR*_*n*_ value (*case-to-normal ratio)* is the ratio of the expression level of gene *n* in the sample under investigation to the average expression level in the control samples. A positive PAS value indicates activation of a pathway, and a negative value indicates repression.

### Building pathway activation profiles and assessment of batch effects

To identify if the OncoFinder technique may improve gene expression analysis by eliminating batch effects, we profiled a set of human clinical bladder cancer tissue samples using the same experimental platform (Illumina human HT 12 v4 bead arrays) in two different laboratories. We investigated gene expression profiles generated from 17 bladder cancer samples and seven normal bladder tissue samples. Eight cancer and four normal samples were analyzed in Dr. Kovalchuk’s laboratory in Lethbridge (Canada), and nine cancer and three normal bladder tissue samples were analyzed in Dr. Buzdin’s laboratory in Moscow (Russia). The gene expression data were deposited in the GEO database (http://www.ncbi.nlm.nih.gov/geo/) with accession numbers GSE52519 and GSE65635.

In agreement with previous reports [Lazar, 2013], the normalized gene expression showed significant batch effects with data from different laboratories clearly clustered on a Principal Component Analysis (PCA) plot (Fig.1A). However, the *PAS* data formed a single merged cluster (Fig.1B). The principle component variability was 4-6 times smaller for the *PAS* data (Fig.1A,B).

**Figure 1.**
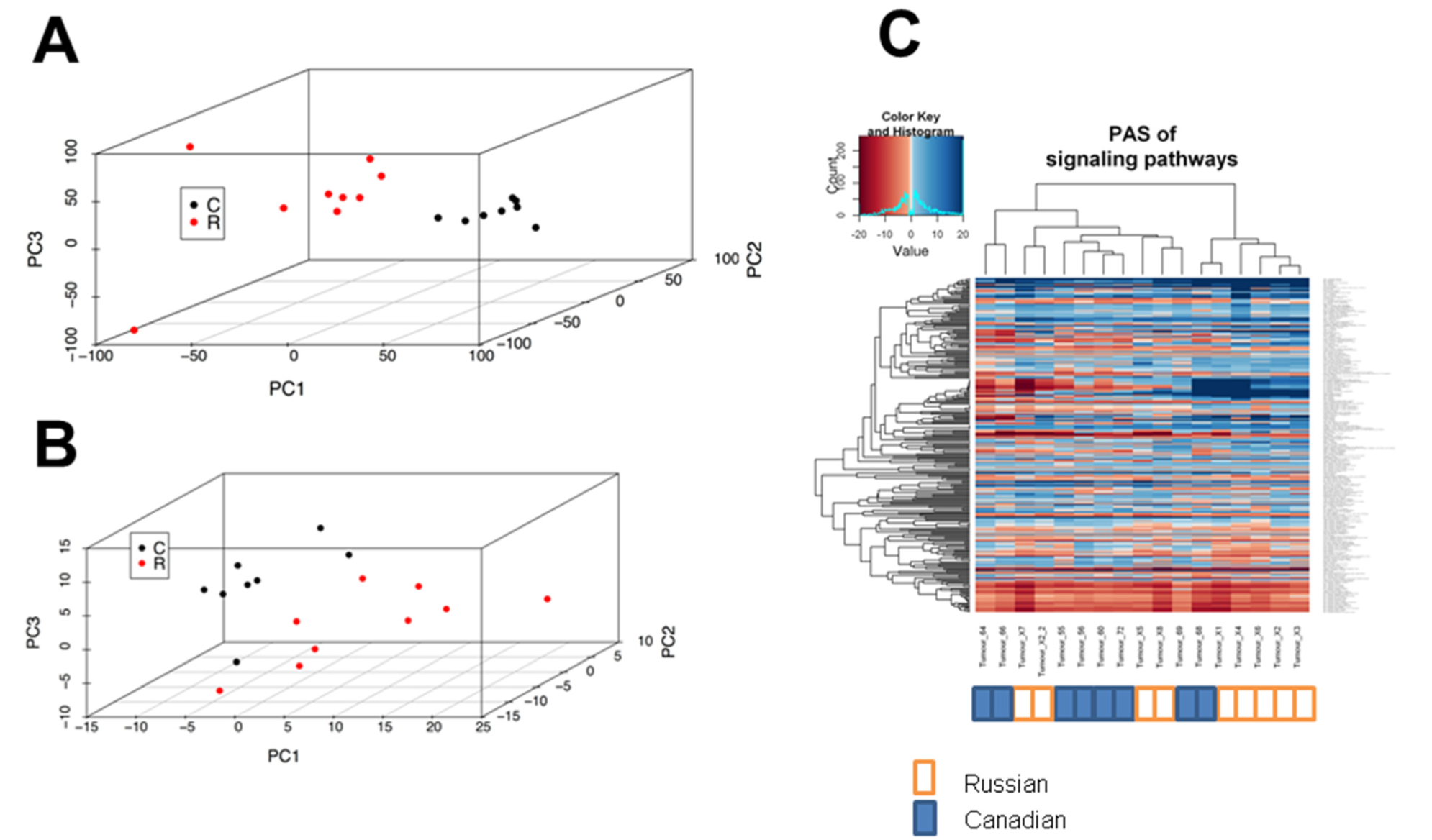
Renal carcinoma datasets assessed at the level of individual gene expression and pathway activation. *A*, principal component analysis (PCA) plot for transcriptomes from datasets obtained in Russia (red dots) and Canada (black dots), at the level of individual gene expression. *B*, PCA plot at the level of molecular pathway activation. *C*, hierarchical clustering dendrogram of the datasets obtained in Russia (marked white) and Canada (marked blue), at the level of molecular pathway activation.

Similarly, using *PAS* values these two sets of samples formed mixed groups on a hierarchical cluster heatmap (Fig.1C). The Canadian samples were labeled 55 - 72; the Russian samples X1 - X8. Some sub-clusters are evidently formed by the samples coming from the different sets, e.g. by samples X5, X8, 69, 68 and X1. (Fig.1C). These data show that data aggregation at the *PAS* level is sufficient to suppress the batch effect in gene expression comparisons.

### Mathematical modeling of data aggregation effects

We investigated the hypothesis that the apparently higher robustness of OncoFinder *PAS* scoring compared to single gene expression, is due to the cumulative nature of the former. *PAS* is the sum of multiple mathematical terms that correspond to each individual gene product participating in a pathway. Model calculations showed that this cumulative effect is able to reduce stochastic noise.

In the model, we included 271 pathways with variable numbers of gene products. We assumed that the expression level of every gene product could be measured using two different methods, say X and Y, corresponding to different experimental platforms (e.g. MH and NGS). Each method introduces errors into the determination of gene expression level, and these errors are independent. A Monte Carlo trial was performed as follows: we simulated both *biased CNR* (with a median value of 1.5) and *unbiased CNR* with a median value of 1. We explored both *noisy* and *exact* expression profiling methods, to allow whether measurement procedures introduce errors in the *true* expression values. The four scenarios of the stochastic simulations (labeled A to D) are shown in Table 1.

**Table 1.**
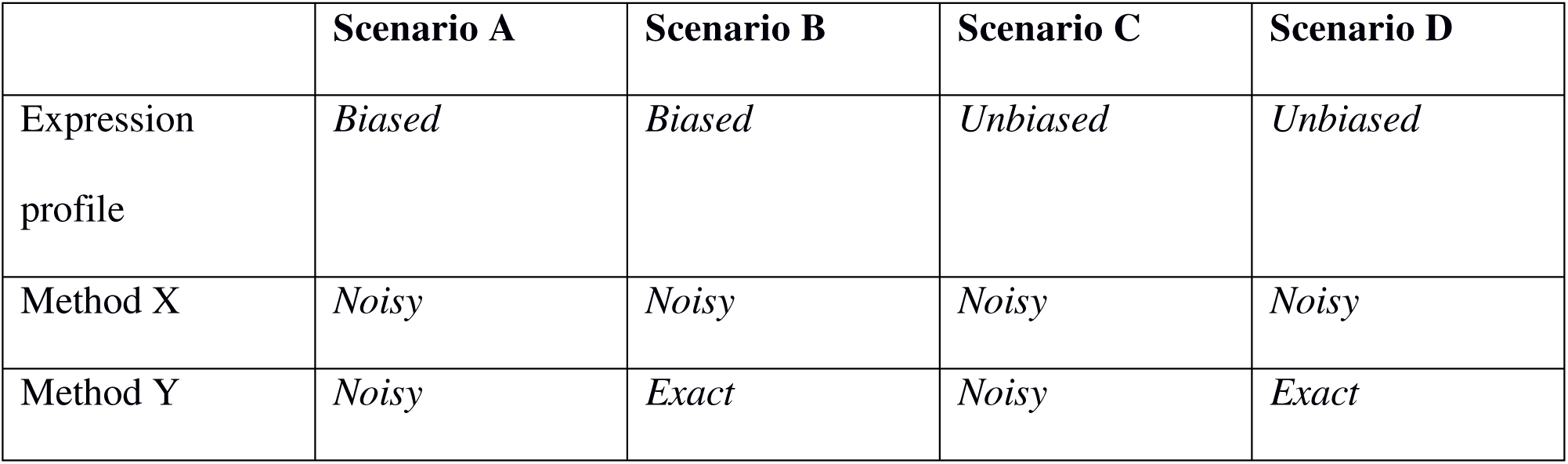
Cross-platform comparisons for modeling the data aggregation effect.

|  | Scenario A | Scenario B | Scenario C | Scenario D |
| --- | --- | --- | --- | --- |
| Expression<br>profile | *Biased* | *Biased* | *Unbiased* | *Unbiased* |
| Method X | *Noisy* | *Noisy* | *Noisy* | *Noisy* |
| Method Y | *Noisy* | *Exact* | *Noisy* | *Exact* |

For each scenario, we calculated the benefit ratio 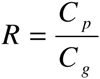, where *C*_*p*_ and *C*_*g*_ are the correlation coefficients between the results obtained using methods X and Y, using *pathway-*based (*PAS*), and *individual* gene product*-*based log *CNR* values, respectively. For each subset of genes in a pathway, we performed 100 Monte Carlo stochastic simulations and then computed the mean values of *C*_*p*_ and *C*_*g*_ using the R statistical package. The greater *R>1*, the higher the benefit from using *PAS* instead of individual gene expression for the cross-platform comparisons; *R<1* means operating at the individual gene product level is better than the *PAS* level.

For *biased* expression profiles, scenarios A and B of Table 1, (Fig. 2), the *PAS* method shows much better agreement between the results obtained using different methods, compared to the individual gene expression levels. The data aggregation advantage of *PAS* is especially strong when both expression methods are *noisy* (scenario A). In scenario B, when one method is *exact*, the benefit of pathway data aggregation is lower. This is caused mainly by higher expression correlation already at the level of individual gene products (Fig. 3). However, the advantages of *PAS* remain considerable for pathways that contain at least 10 gene products (Fig. 2). For shorter pathways, the data aggregation effect is gradually decreased, and the *R* ratio reflecting the benefit of using *PAS* values, trends towards 1.

**Figure 2.**
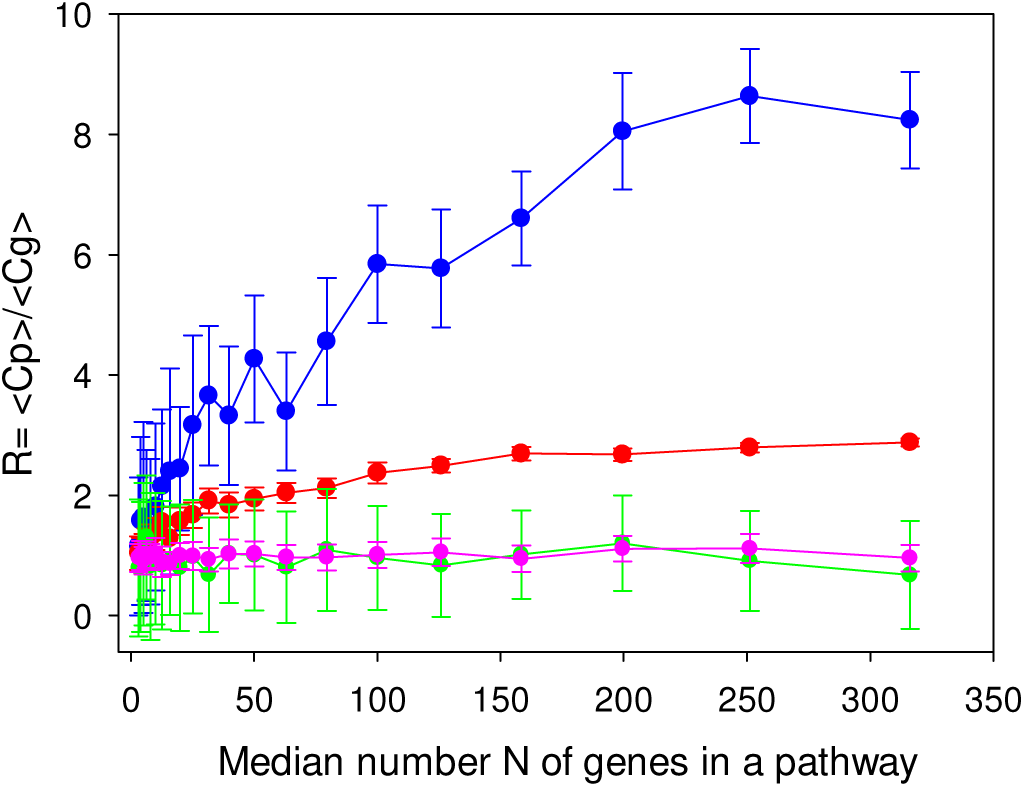
Ratio of pathway-related and gene-related correlation coefficients between results obtained using hypothetical methods X and Y, as a function of the median gene number, N, in a pathway for four scenarios: A (blue) – *biased* expression profile, *noisy* method Y; B (red) - *biased* expression profile, *exact* method Y; C (green) – *unbiased* expression profile, *noisy* method Y; D (magenta) – *unbiased* expression *exact* method Y. The method X is always condsidered *noisy*.

**Figure 3.**
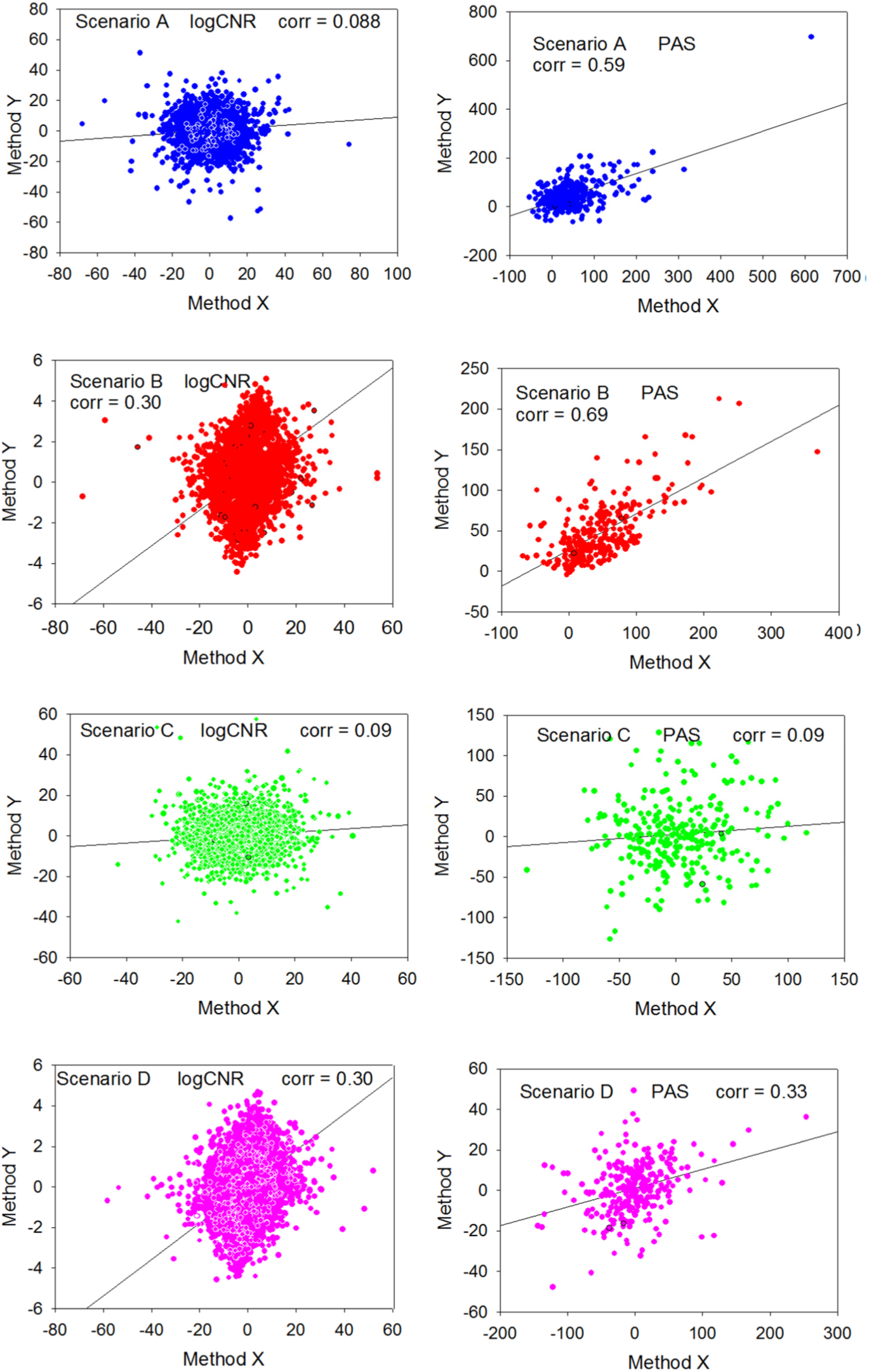
Distributions of values obtained during random trials using two different expression profiling methods X (horizontal axis) and Y (vertical axis). Median number of gene products in a pathway is 100. Left column: log *CNR* for individual gene products, method Y vs method X. Right column: *PAS* scoring method Y vs method X. Blue dots: scenario A (*biased* expression profile, *noisy* method Y). Red dots: scenario B (*biased* expression profile, *exact* method Y). Green dots: scenario C (*unbiased* expression profile, *noisy* method Y). Magenta dots: scenario D (*unbiased* expression profile, *exact* method Y). Method X is always considered *noisy*.

For *unbiased* transcription profiles, with median relative gene expression levels equal to 1, the data aggregation effect is completely lost (scenarios C and D). Here, the mean value for each gene product component of the *PAS* score is zero; consequently, the expected *PAS* is also zero, and the relative data variation is the same at the gene product and the *PAS* level.

The simulations clearly elucidate how the cumulative nature of *PAS* suppresses cross-platform data variation and batch effects. They show that there is a significant advantage of using *PAS* to compare platforms, when at least one is *noisy*. This should apply to most if not all existing high throughput experimental platforms, and it should be seen when experimental expression data is compared. The simulations demonstrate that *PAS* calculations are advantageous for *biased* transcriptomes and proteomes and virtually useless for *unbiased* ones. Unbiased data sets are too similar to the control group used as the reference to calculate *CNR* values. This means that the *PAS* approach will be especially useful when the expression signature in the sample under study is very different from that of the control samples. This finding may help to identify appropriate control samples for decreasing expression data noise. Finally, this model shows that the higher is the number of gene products in a pathway, the greater the benefit of shifting from individual gene/protein expression to *PAS* data For example, the mean number of gene products in the OncoFinder database is 68 per pathway, and the model predicts about a 4.5 –fold decrease in data variation at the *PAS* level in the biased noisy-noisy scenario, which may explain the success of the OncoFinder approach in various applications [Buzdin, 2014a].

### Experimental model of cross-platform comparisons

In transcriptomic methods, batch effects arise from errors introduced at the stages of RNA purification, library preparation and amplification, hybridization and reading of arrays [Risso, 2011]. We investigated whether the OncoFinder *PAS* algorithm can suppress batch effects introduced by cross-platform comparisons. At the same time we assessed if the algorithm works efficiently for formalin-fixed, paraffin-embedded (FFPE) tissue samples. Seven FFPE tissue blocks isolated from human renal carcinomas were profiled using two independent experimental platforms. The first was the Illumina HT 12 v4 bead array system optimized for FFPE tissues. The second was a customized microchip system developed using the CustomArray (USA) technology of direct on-chip electrochemical oligonucleotide synthesis. The custom arrays had 3775 oligonucleotide probes corresponding to 2214 human gene products involved in 271 intracellular signaling pathways (Supplementary dataset S1). The custom arrays, used the original oligonucleotide probe sequences of the Illumina HT 12 v4 platform, but shortened by 5 nucleotides at the 5’ end and by 5 nucleotides at the 3’ end. Quantile –normalized gene expression data were deposited into the GEO database with the accession numbers GSE65637 and GSE65639. The differences between the Illumina and the Custom platforms included shorter oligonucleotide probe sequences, different library preparation protocols and different hybridization signal development and reading methods (Supplementary Fig.1). The Custom method for library preparation was quite distinct from Illumina and identical to that used by the Agilent MH platform (Supplementary Fig.1B,C,E) with the sole exception that biotinylated rather than fluorescently labeled DNA is used at the terminal stage (Supplementary Fig.1 B,E). A brief comparison of the protocols used for the Custom and top commercial MH platforms manufactured by the Illumina, Agilent and Affymetrix companies for FFPE tissue profiling is given in Supplementary dataset S2.

To compare with the renal carcinoma samples, we used GEO dataset GSE49972 [Karlsson, 2014] containing 6 normal kidney samples to normalize the expression data and calculate *PAS*. The normalized *CNR* expression data and *PAS* values are shown in Supplementary dataset S1. At the level of individual gene products, we observed relatively low correlations (0.2-0.3) between the same transcriptomes profiled using the two platforms (Fig. 4; Supplementary dataset S3). In contrast, at the *PAS* level the correlations were strong, varying from 0.84 to 0.91 (Fig. 4; Supplementary dataset S3).

**Figure 4.**
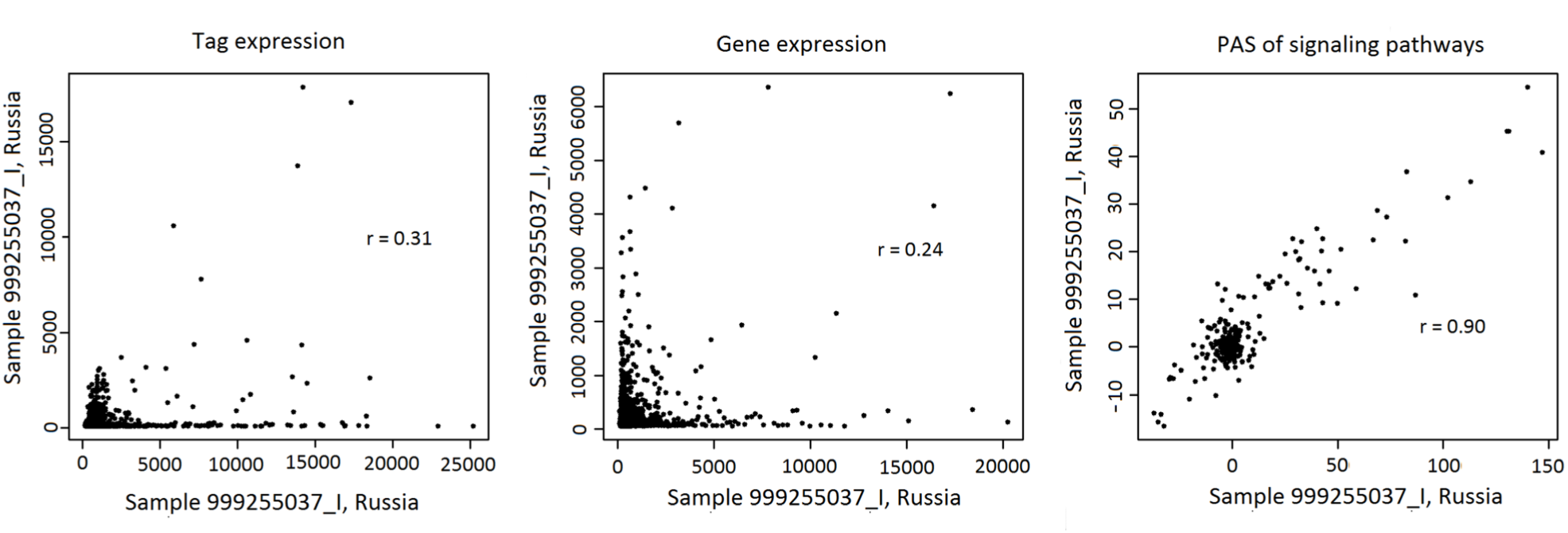
Correlation between transcriptomic data obtained for the same representative renal carcinoma specimen using the Illumina HT12 (ordinate) and CustomArray (abscissa) microarray platforms. The panels represent (from left to right) correlation between the oligonucleotide expression tags, correlations at the level of individual genes, and correlation at the level of molecular pathways.

These results experimentally confirm the hypothesis that data aggregation at the *PAS* level increases the stability of cross-platform expression data and that the advantage of *PAS* is retained for FFPE samples.

### Data aggregation effects assessed on different RNA and protein expression profiles

We investigated quantitative aspects of the effect of data aggregation on several datasets where the same samples were profiled using different expression platforms (Tab. 2, Supplementary dataset S4).

**Table 2.** Transcriptomic and proteomic datasets used to assess data aggregation effects.

| **Paper reference, Dataset ID** | **Origin** | **Case and control samples** | **Experimental platforms** | **Number of samples** |
| --- | --- | --- | --- | --- |
| [van Delft, 2012],<br>GSE36244 | HepG2 cells | Cells treated with benzopyrene (*cases*) vs untreated cells (*norms*) | Transcriptomes using Affymetrix Human Genome U133 Plus 2.0 arrays and Illumina Genome Analyzer sequencer | 4 |
| [Xu, 2013],<br>GSE41588 | HT-29 cells | Cells treated with 5-aza-deoxy-cytidine (*cases*) vs untreated cells (*norms*) | Transcriptomes using Affymetrix Human Genome U133 Plus 2.0 arrays and Illumina Genome Analyzer sequencer | 6 |
| [Kim, 2013]<br>GSE37765 | Lung adeno-carcinoma | Tumor samples (*cases*) vs normal lungs (*norms*) | Transcriptomes using Agilent 1M CNV arrays and Illumina Genome Analyzer sequencer | 6 |
| This study | Renal carcinoma tissue | Tumor samples (*cases*) vs normal adult kidneys (*norms*) | Transcriptomes using Illumina Human HT-12 v4 microarrays and Custom microchip platform (see text) | 7 |
| [Yang, 2014],<br>GSE52488,<br>PXD000624 | Human smooth muscle cells | Cells treated with PDGF served as *cases*, untreated - as *norms*. | Transcriptome using Affymetrix Human Gene 1.0 ST arrays and proteome using triplex SILAC at Orbitrap XL mass spectrometer. | 2 |
| [Cabezas-Wallscheid, 2014], EMTAB-2262, PXD000572 | Murine hemato-poietic stem cells (HSC) | HSC served as *norms*, multipotent progenitor population 1 (MPP1) – as *cases*. | Transcriptome using RNA-seq HiSeq2000 (Illumina) and proteome using duplex SILAC at Orbitrap Velos Pro mass spectrometer | 4 |
| [Hara, 2013] | Human pathologic skin fibroblasts | Samples from two patients served as *cases*. Three and two normal samples were used as *norms* for proteome and transcriptome investigation, respectively | Transcriptome using Affymetrix Human Genome U133 Plus 2.0 arrays and proteome using triplex SILAC at Orbitrap Velos mass spectrometer | 2 |

We observed two trends for the behavior of the benefit ratio 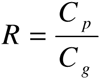. In model calculations, we observed a crucial role of expression profile bias between the *case* and *normal* samples for successful data aggregation of genes into pathways (Fig. 2, 3). We introduce a measure of such bias, termed 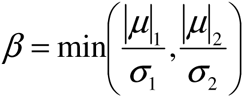, where ***μ***_*i*_ and ***σ***_*i*_ are the mean and standard deviation, respectively, of the set of log *CNR* values obtained for a given sample using the experimental platform *i*. The results of the model calculation (Fig. 2,3, scenarios A and B) suggest that, even for the same values of *β*, *R* may be different depending on *C*_*g*_ (correlation at the individual gene product level): the higher *C*_*g*_, the lower *R* at equal *β*.

With a discrimination threshold for *C*_*g*_ chosen as equal to 0.25 between *low-correlated* and the *considerably correlated* samples, we can see the clear clusters of data for data aggregation effect (Fig. 5, blue dots for *low* and red dots for *considerably* correlated samples. Note that the two clusters of data depending on the *C*_*g*_ threshold are seen for both transcriptome-to-transcriptome and transcriptome-to-proteome comparisons.

**Figure 5.**
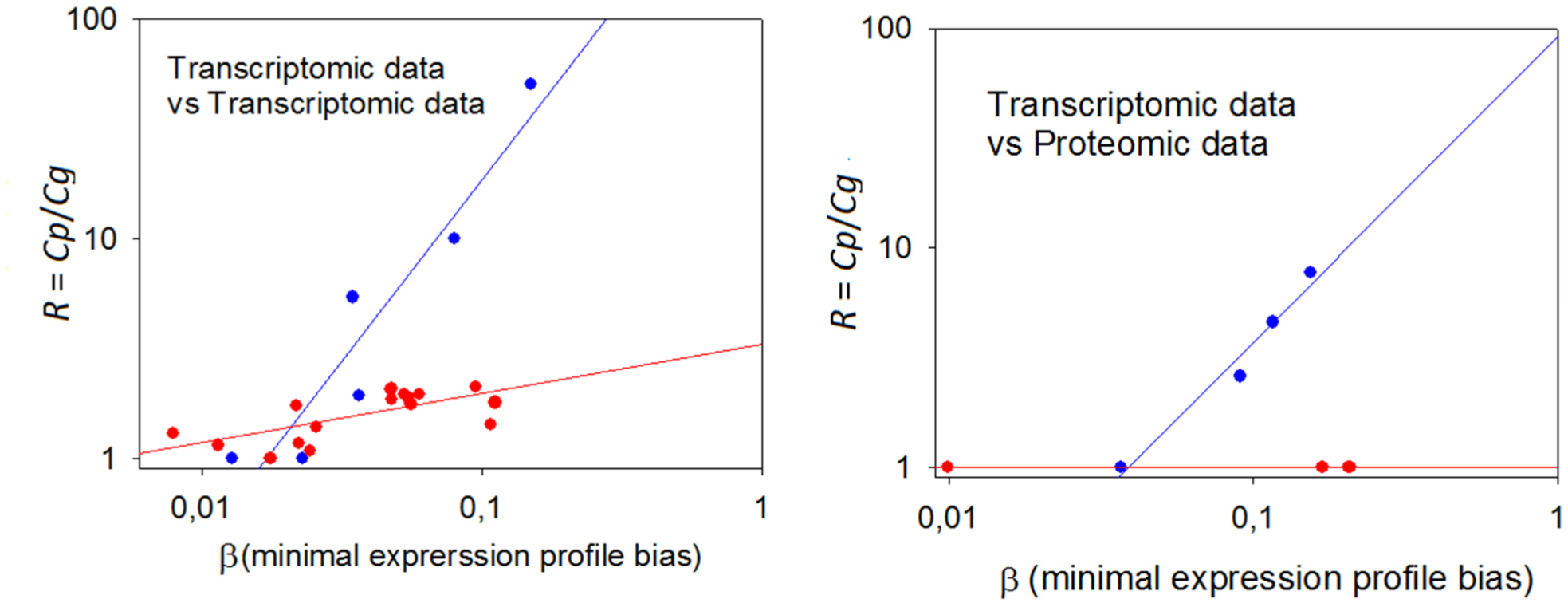
Dependence of the data aggregation effect (*R*) on the minimal expression profile bias *β*. Left panel: transcriptome-to-transcriptome comparisons for the same samples using different experimental platforms. Right panel: transcriptome-to-proteome comparisons for the same samples. The *C*_*g*_ threshold between the samples *low* and *considerably* correlated at the gp level was chosen as equal to 0.25; blue dots: *low* correlation at gene product level; red dots: *considerable* correlation at gene product level.

The data obtained suggests that when *β* is low, the *R* is hardly distinguishable from 1; however, when *β* exceeds a threshold, the increase of *R* becomes statistically significant. Finally, these results also demonstrate that transcriptomic and proteomic profiles demonstrate more compatible results at the molecular pathway level rather than on the level of individual gene products.

### Comparison of PAS scoring methods according to their capacities in data aggregation

We compared the abilities of five popular *PAS* scoring methods to yield an advantageous the data aggregation effect when the expression of molecular pathways is compared instead of individual gene products. For the seven renal carcinoma samples discussed above, we calculated *R* using alternative *PAS* scoring methods: OncoFinder [Buzdin, 2014b], *topology analysis of pathway phenotype association*, TAPPA [Gao, 2007], *topology-based score* (TB) [Ibrahim, 2012], *pathway-express* (PE) [Draghici, 2007], and *signaling pathway impact analysis* (SPIA) [Tarca, 2009] methods (Supplementary data set S5). These methods differ in the factors used to evaluate the importance of distinct gene products in pathway activation.

Only three of the methods, OncoFinder, PE and SPIA, showed a substantial data aggregation effect (*R*) ranging from 2-2.3. Other methods showed lack of any positive effect (Fig. 6).

**Figure 6.**
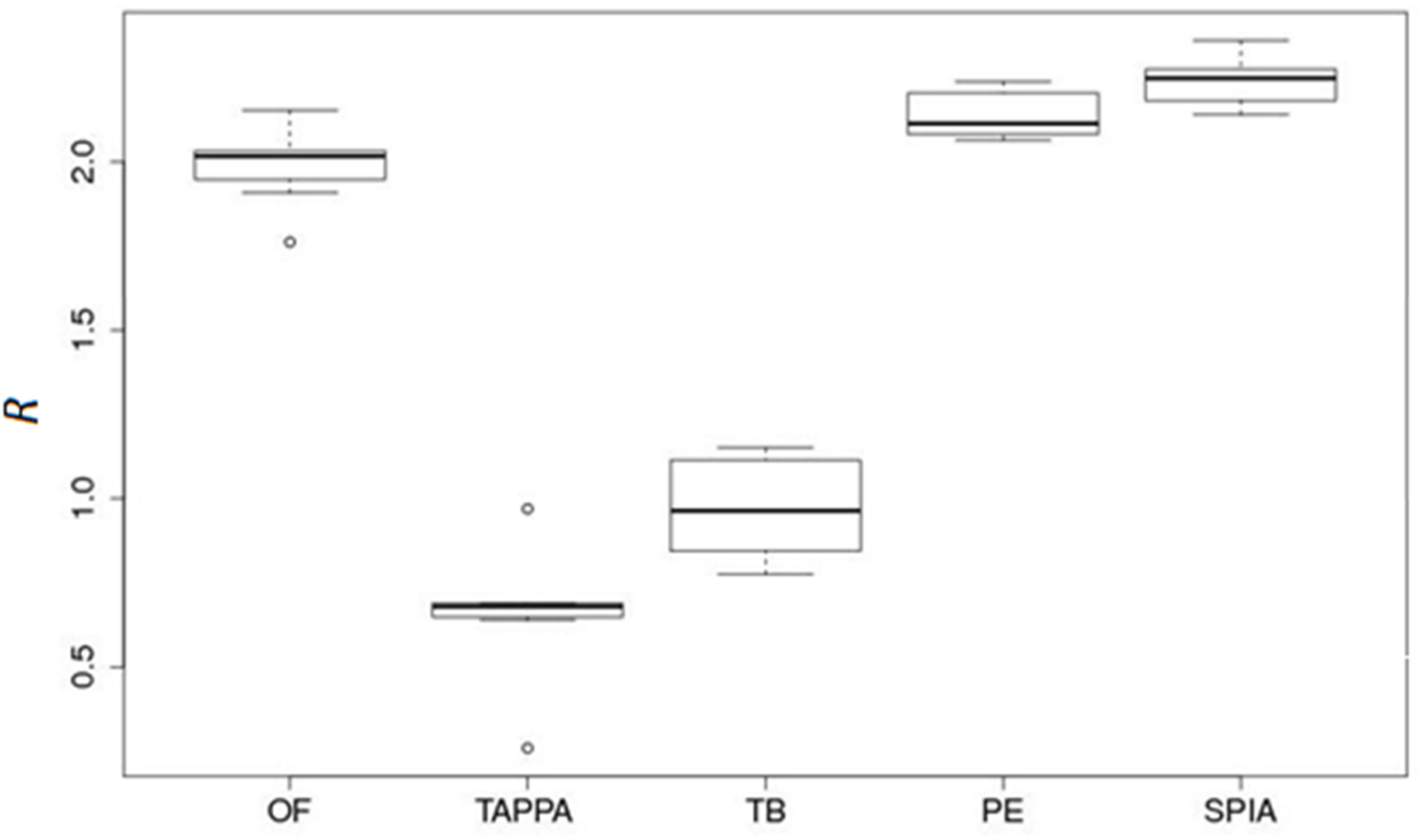
Data aggregation effect *R* for five pathway activation scoring methods (OncoFinder, TAPPA, TBScore (TB), Pathway-Express (PE), and SPIA) on the renal carcinoma dataset.

### Different methods for PAS scoring show different properties in retention of biological features

Cross-platform data comparison has the potential to become an extremely useful tool in contemporary biomedicine and bioinformatics. Although the application of *PAS* methods has the ability to restore correlations between different expression data sets, the absolute values of *PAS* may differ between platforms. To overcome this inconsistency, several *cross-platform harmonization*^1^ methods can be applied ranging from the simplest z-scaling and mean-centering to more sophisticated algorithms utilizing machine-learning/Bayesian harmonization (e.g., [Warnat, 2005; Shabalin, 2008; Hsu, 2014], including the popular harmonization technique XPN [Shabalin, 2008]. In many applications these harmonization methods can diminish the systematic bias introduced by the experimental methods and devices used, but they demonstrate lower efficiencies for routine batch effects like those observed when comparing results obtained using the same platform but on different calendar dates or in different laboratories.

This made it of interest to compare the ability of the five *PAS* scoring methods to retain biological features after cross-platform data harmonization with the XPN method.

We used the results of the Microarray quality control project (MAQC) [MAQC Consortium, 2006] as a model dataset for this study. The MAQC project investigated four types of samples (A-D; each sample profiled in 15 technical replicates) using different microarray devices. Type A samples were taken from the Stratagene Universal Human Reference RNA; type B samples – from the Ambion Human Brain Reference RNA. Type C and D samples were obtained by combining samples A and B in mass ratios 75:25 for C, and 25:75 for D, respectively.

After XPN harmonization of gene expression profiling using the Agilent Whole Human Genome Oligo and Affymetrix Human Genome U133 Plus 2.0 platforms, we applied different methods of *PAS* scoring (Supplementary dataset S6) using the samples of type A as *normal*. The probability densities of the Euclidean distances between the *PAS* vectors calculated for the three samples (B, C, and D) differ greatly depending on the *PAS* scoring method used (Fig.7). In such an assay, an ideal *PAS* scoring method should make distinctions between samples depending primarily on the sample types, rather than on the experimental platform used. A satisfactory *PAS* calculation method, therefore, should yield a unimodal distribution of the *PAS-PAS* distances, without any significant deviations. If the distribution of *PAS-PAS* distances is bimodal or multimodal, this points to the inability to eliminate platform-specific bias even at the pathway level. Only the OncoFinder and TAPPA methods were able to eliminate the cross-platform bias for all three sample types (Fig.7).

**Figure 7.**
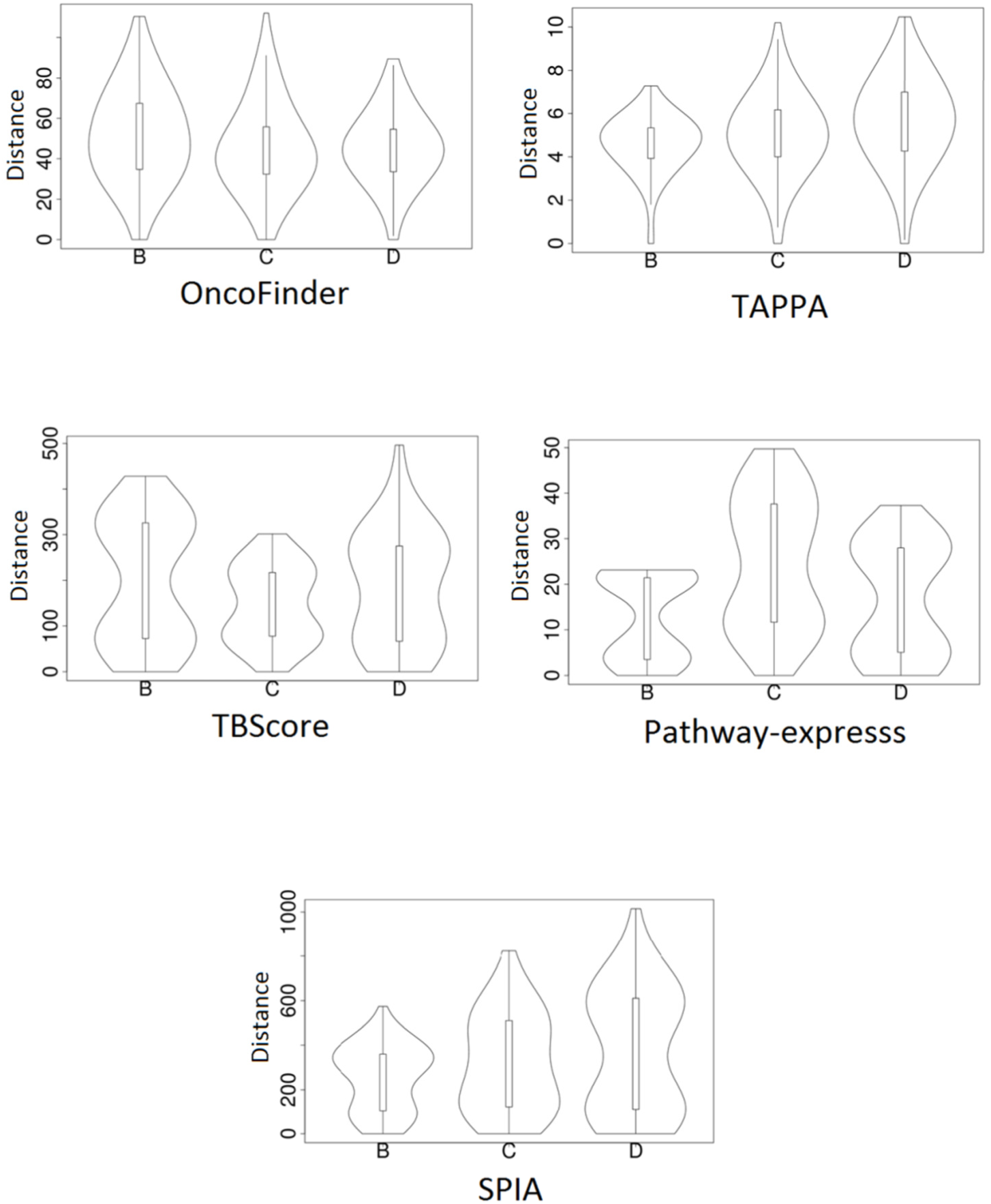
Distribution of Euclidean distances between the *PAS* vectors for different sample types taken from the MAQC dataset (marked as B, C and D) using different methods of *PAS* scoring. A *unimodal* distribution indicates lack of significant difference between within-platform and cross-platform distances. A *bimodal* distribution means that the cross-platform *PAS* distance (upper mode in the violin plots) is essentially higher that the within-platform distance. See text for descriptions of the different scoring methods.

Hierarchical clustering (dendrograms shown in Supplementary data set S7). demonstrates that only the OncoFinder and TAPPA methods enabled clustering of the *PAS* vectors exclusively according to biological sample type. Thus, among the five *PAS* scoring algorithms tested, only OncoFinder showed effective data aggregation with efficient retention of biological information in three independent tests (Table 3).

**Table 3.** Comparison of *PAS* scoring methods using functional and statistical tests.

| **Method** | **Data aggregation**<br>**effect** | **Distance distribution**<br>**within each sample type** | **Quality of *PAS***<br>**clustering** |
| --- | --- | --- | --- |
| OncoFinder | ++ | +++ | +++ |
| TAPPA | -- | +++ | ++ |
| TBScore | - | -- | - |
| Pathway-express | +++ | -- | -- |
| SPIA | +++ | -- | -- |

## DISCUSSION

High throughput gene expression may produce both random and systematic errors, arising from the steps in RNA or protein purification, library preparation and/or amplification, hybridization and sequencing, reading arrays, and mapping and annotation of the reads [Chalaya, 2004; Shugay, 2014; Risso, 2011]. It is generally hard to identify the types of errors and to find out which kind of experimental protocol provides more reliable data. While the measured concentration of each individual gene product may be in error, we show in this report that combining sufficient numbers of these concentrations into a pathway-oriented network apparently generates significantly more stable data. We also tested whether OncoFinder and other *PAS* scoring methods can improve expression data to suppress batch effects, the unwanted variation in gene expression measurements on the same experimental platform made at different times, which frequently originate from the limitation in the number of samples that can be processed at once in a single experiment [Demetrashvili, 2010]. Batch effects also hinder the combination of different experimental datasets. Batch effects are almost inevitable [Lazar, 2012]. By limiting analyses to single data sets, one frequently must use an insufficient number of samples, which leads to high false-negative rates [Lazar, 2012]. Eliminating batch effects enables larger datasets, and provides more statistical power to subsequent analyses [Lazar, 2012].

Here, using the Illumina HT12 bead array platform to profile human cancer samples, we demonstrate that the *PAS* scoring technology OncoFinder effectively suppresses batch effects present in the individual gene expression measurements (Fig.1). OncoFinder efficiently increases expression data stability from all major experimental platforms, for both fresh and formalin-fixed, paraffin-embedded tissue samples (Fig.4).

Various publicly available repositories of gene expression data embrace the full spectrum of normal and pathological conditions for the majority of known human diseases [Cancer Genome Atlas Research Network, 2008; Jones, 2006]. Unfortunately, batch effects, which bias the expression profiles, hamper the joint analysis of most of this data obtained using different experimental settings.

Discrepancies in data obtained on *the same* and *different* experimental platforms, rmust be addressed by different methods, termed *normalization* and *harmonization*, respectively. For intra-platform normalization, more attention is paid to equilibration of scaling factors, while cross-platform harmonization must address the type of distribution of output intensities for each gene. Exiting methods for intra-platform normalization include quantile normalization [Bolstad, 2003] and frozen robust multi-array analysis (FRMA) [McCall. 2010] for microarray data, and the DESeq method [Anders, 2010] for next-generation sequencing.

Methods for cross-platform harmonization, such as distance-weighted discrimination (DWD) [Huang. 2012], cross-platform normalization (XPN) [Shabalin, 2008], and platform-independent latent Dirichlet allocation (PLIDA) [Deshwar, 2014], provide deep restructuring or signal intensity redistribution for the entire set of genes profiled. As a rule, the cross-platform harmonization involves data clustering and finding similarity regions among results obtained using different platforms, to strengthen similarity during the harmonization process.

Unfortunately, current normalization and harmonization methods hardly distinguish between artifacts introduced by batch effects and the real biological differences. Additional tools are needed to improve normalization and harmonization procedures. We demonstrate here for most major transcriptomic and proteomic commercial platforms that data aggregation at the level of molecular pathways has the potential to reduce greatly the bias in the datasets under comparison. Since each pathway may contain hundreds of different gene products, transition from single gene products to the whole pathway level may restore biologically significant correlations.

We propose a term *data aggregation effect* for such restoration of biological correlation at the pathway level. We created a mathematical model that simulates it and identifies the necessary conditions for its applicability. Sample expression profiles must be biased compared to control samples, i.e. the transcriptional signatures of the *case* samples must differ significantly from the normal ones (Fig. 5). The strength of the data aggregation effect grows with the number of gene products in a molecular pathway. The data aggregation effect is especially strong when the initial correlation between the expression data is weak (Fig. 2,3). Finally, the choice of *PAS* scoring method affects the data aggregation effect. On a model data set, the OncoFinder, Pathway-Express and SPIA algorithms result in a considerable data aggregation effect, while TAPPA and TB-Score don’t (Fig. 6). Only OncoFinder and TAPPA were able to preserve the biological features on the model dataset MAQC after cross-platform harmonization, while with Pathway-Express, SPIA and TB-Score methods, platform-introduced bias features still dominated the output expression signatures (Fig.7). Thus, among the five *PAS* scoring methods tested here, the OncoFinder algorithm showed the best efficiency and accuracy (Tab.3), which makes OncoFinder a method of choice for many applications using high-throughput analysis of gene expression at the RNA or protein levels.

It should be possible in the future to refine *PAS* methods to create universal platform-agnostic analytic tools. These tools have a huge potential to accelerate progress in genetics, physiology, biomedicine, molecular diagnostics and other applications by combining unbiased data from many sources and various experimental platforms.

## MATERIALS AND METHODS

### Tissue collection and RNA isolation from fresh biosamples

Seven normal bladder and seventeen bladder carcinoma specimens from patients treated at the P.A. Herzen Moscow Oncological Research Institute (HMORI; Moscow, Russia) were analyzed. Of these samples (cancer/normal), nine/three were examined at the Shemyakin-Ovchinnikov Institute of Bioorganic Chemistry (IBC; Moscow, Russia) and eight/four at the University of Lethbridge (UL; Alberta, Canada). All patients provided written informed consent to participate in this study. This study was approved by the local ethical committees at IBC, UL and HMORI. Tumor samples were obtained from patients who had undergone surgery for bladder carcinoma at the HMORI between 2009 and 2013. The median age of the cancer patients at the time of surgical tumor resection was 64 years (range 48–77 years). Tissue samples from non-cancer controls were collected from autopsies at the Department of Pathology at the Faculty of Medicine, Moscow State University. Both the tumors and normal tissues were evaluated by a pathologist to confirm the diagnosis and estimate the tumor cell numbers. All tumor samples used in this study contained at least 80% tumor cells. The median age of the healthy tissue donors was 45 years (range 20–71 years). Tissue samples were stabilized in RNAlater (Qiagen, Germany) and then stored at −80°C. Frozen tissue was homogenized in TRIzol Reagent (Life Technologies, USA), and RNA was isolated following the manufacturer’s protocol. Purified RNA was dissolved in RNase-free water and stored at −80°C.

### Microarray profiling of gene expression in fresh biosamples

Total RNA was extracted using TRIzol Reagent and then reverse-transcribed to cDNA and cRNA using the Ambion TotalPrep cRNA Amplification Kit (Invitrogen, USA). The cRNA concentration was quantified and adjusted to 150 ng/ml using an ND-1000 Spectrophotometer (NanoDrop Technologies, USA). 750 ng of each RNA library was hybridized onto the bead arrays.

Gene expression experiments were performed by Genoanalytica (Moscow, Russia) and the O. Kovalchuk Laboratory (Lethbridge, Canada) using the Illumina HumanHT-12v4 Expression BeadChip (Illumina, Inc.). This gene expression platform contains more than 25,000 annotated genes and more than 48,000 probes derived from the National Center for Biotechnology Information RefSeq (build 36.2, release 22) and the UniGene (build 199) databases. The expression data were deposited in the GEO database (http://www.ncbi.nlm.nih.gov/geo/), accession numbers GSE52519 and GSE65635.

### Synthesis of microarrays

A B3 synthesizer (CustomArray, USA) was used for oligonucleotide probe synthesis on the CustomArray ECD 4X2K/12K slides. Synthesis was performed according to the manufacturer’s recommendations. At least three replicates of total 3823 unique oligonucleotide probes of 40 nucleotides in length for 2278 genes were placed on each chip.

### Library preparation and hybridization

RNA was extracted from freshly frozen tissue samples or samples stored in stabilizing buffer solutions using the standard protocol for TRIzol reagent (Life Technologies). RNA extraction from FFPE samples was performed using the RecoverAll™ Total Nucleic Acid Isolation Kit for FFPE. Complete Whole Transcriptome Amplification WTA2 Kit (Sigma) was used for reverse transcription and library amplification. The manufacturer’s protocol was modified by adding to amplification reaction a dNTP mix containing biotinylated dUTP, resulting to a final proportion dTTP/biotin-dUTP of 5:1.

Hybridization was performed according to the CustomArray ElectraSense™ Hybridization and Detection protocol. The hybridization mix contained 2.5 ug of labeled DNA library, 6X SSPE, 0.05% Tween-20, 20mM EDTA, 5x Denhardt solution, 100 ng/ul sonicated calf thymus gDNA, and 0,05% SDS. The chip was incubated in the hybridization mix overnight at 50ºC. The hybridization efficiency was detected electrochemically using CustomArray ElectraSense™ Detection Kit and ElectraSense™ 4X2K/12K Reader. The chip was designed using the Layout Designer software (CustomArray, USA).

### Functional annotation of gene expression data

The SABiosciences (http://www.sabiosciences.com/pathwaycentral.php) signaling pathways knowledge base was used to determine structures of intracellular pathways, as described previously [Spirin, 2014].

*OncoFinder.* We applied the original OncoFinder algorithm [Buzdin, 2014b] for functional annotation of the primary expression data and for calculating *PAS* scores. The microarray gene expression data were quantile normalized according to [Bolstad, 2003]. The formula used to calculate the *PAS* for a given sample and a given pathway *p* is as follows:

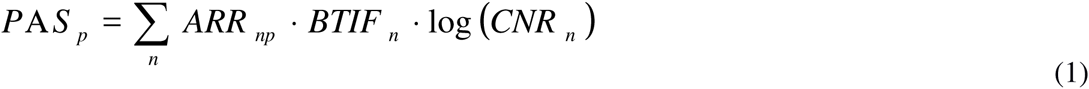

Here the case-to-normal ratio, *CNR*_*n*_, is the ratio of the expression level of gene *n* in the sample under investigation to the average expression level of that gene in the control group of samples. The Boolean flag of *BTIF* (beyond tolerance interval flag) equals one or zero when the *CNR* value has simultaneously passed or not passed, respectively, the two criteria that indicate a significantly perturbed expression level from an essentially normal expression level. The first criterion is that the expression level of the sample lies within the tolerance interval, with *p* < 0.05. The second criterion is whether the *CNR* value lies outside the cut-off limits, i.e., either *CNR* < 2/3 or *CNR* > 3/2. *ARR*_*np*_, the discrete value of the activator/repressor role equals the following fixed values: −1, when the gene/protein *n* is a repressor of molecular pathway; 1, if the gene/protein *n* is an activator of pathway; 0, when the gene/protein *n* is known to be both an activator and a repressor of the pathway; and 0.5 and −0.5, respectively, tends to be an activator or a repressor of the pathway *p*, respectively.

Our approach to calculations of *PAS* implies two principal assumptions:

1) First, computational modeling of signal transduction processes [Birtwistle, 2007; Borisov, 2009; Kuzmina, 2011] indicates that for most interacting proteins the concentration of their active forms, which are sufficient for downstream signaling, is much lower than the total abundance of the corresponding protein. In other words, signal transduction may be performed even at the very low level for most gene products.

2) Second, we stipulate that each pathway graph may be simplified up to the following structure that includes only two chain-like (linear) branches: one for sequential events that promote activation of whole pathway, and another for repressor sequential events. The adequacy of this quite radical approximation was shown before in comparison with the full-scaled kinetic model [Kuzmina, 2011], when all protein-protein interactions were described using the mass-action law along each edge of a highly branched pathway graph [Buzdin, 2014].

Under these conditions, we presume that all activator/repressor members have equal importance for the whole pathway, and come to the following formula for the overall signal outcome (*SO*) of a given pathway, 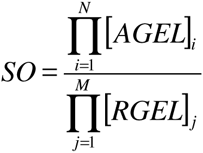. Here the multiplication is done over all possible activator and repressor proteins in the pathway, [*AGEL*]_*i*_ and [*RGEL*]_*j*_ are relative gene expression levels of activator (*i)* and repressor (*j)* members, respectively. To obtain an additive value, it is possible to take the logarithmic levels of gene expression, and thus come to a function of *PAS*.

The results for 271 pathways were obtained for each sample (see Supplementary Data set S1). Statistical tests used the R software package.

*TAPPA (Topology analysis of pathway phenotype association).* Imagine a pathway graph, *G*(*V,E*) where *V* = {*g*_1_,*g*_2_,…,*g*_n_} is the set of graph nodes (vertices), and *E* = {(*g*_i_,*g*_j_) | genes *g*_i_ and *g*_j_ interact} is the set of graph edges [Gao, 2007]. The adjacency matrix is defined as follows, *a*_*ij*_ = 1, if *i* = *j* or (*g*_*i*_,*g*_*j*_)∈*E*, and *a*_*ij*_ = 0, if (*g*_*i*_,*g*_*j*_)∉*E*. A *centered Z-scoring* procedure was applied to the logarithmic gene expression matrix 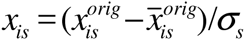.

The adjacency index for a pathway is defined as follows,

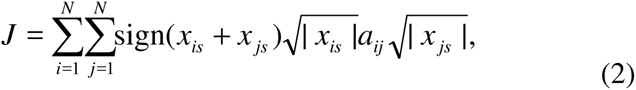

where *N* is the number of genes in the pathway, and the double summation of over the sign(*x*_*is*_ + *x*_*js*_)reveals whether the pathway has more up- or down-regulated genes. The sign (of what?), indicates whether the whole pathway is up- or down-regulated is calculated as 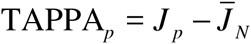, where 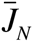 is the expected value of *J* over the set of samples that are considered normal.

*TBScore (Topology-based score)* [Ibrahim, 2012]. For a pathway *p* that has *N* nodes, the value 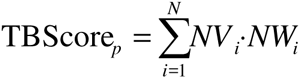, where the node value, *NV*, equals to zero if all the genes in the node *i* are non-differential genes, or equals to the sum of log-fold-changes of the differential genes in the node *i*. The gene is considered differential, if the gene is considered differential in terms of the Boolean flag *BTIF* (as for the OncoFinder algorithm). The node weight, *NW*_*i*_, equals the number of downstream nodes for node *i*. To determine the value of *NW*_*i*_, we used the depth-first search method [Even, Sh. Graph Algorithms, Cambridge University Press, 2011] using labeling visited nodes to avoid the infinite cycling.

*Pathway-Express (PE)* [Draghici, 2007]. The PE-score for a pathway *K* was calculated as follows,

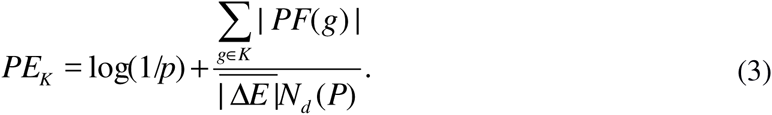

The first term in this sum is the p-value for the probability to obtain the observed or a higher number *N*_*d*_ of differentially expressed genes (between the pools of case and normal samples) by random chance, assuming a hypergeometrical distribution for *N*_*d*_. The second term is a summation over the perturbation factors (*PF*) for the all genes *g* of the pathway *K*,

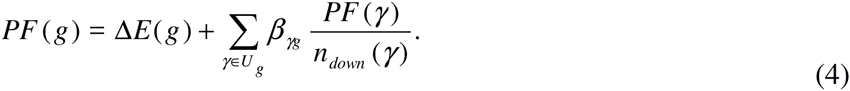

Here *ΔE*(*g*) is the signed difference of gene *g* logarithmic expression in a given sample compared with the expected value for the pool of normal samples. The latter term expresses the summation over all the genes γ that belong to the set *U*_*g*_ of the upstream genes for the gene *g*. The value of *n*_*down*_(γ) denotes the number of downstream genes for gene γ. The weight factor *β*_*γg*_ indicates the interaction type between γ and *g*: *β*_*γg*_ = 1 if γ activates *g*, and *β*_*γg*_ = −1 when γ inhibits *g*.

Although the value of *PF* may be positive or negative, the overall score of *PE* is obligatory positive. The search for upstream/downstream genes is performed according to the depth-first search method, as in the TBScore method.

*SPIA (Signal pathway impact analysis)* [Tarca, 2009]. To obtain an estimator for pathway perturbation that is positive for an up-regulated and negative for a down-regulated pathway, use the second term in formula (4), resulting in the accuracy value, *Acc*(*g*) = *PF*(*g*) − Δ*E*(*g*). It can be shown that [Tarca, 2009] this accuracy vector may be expressed as follows,

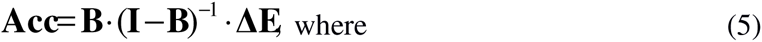

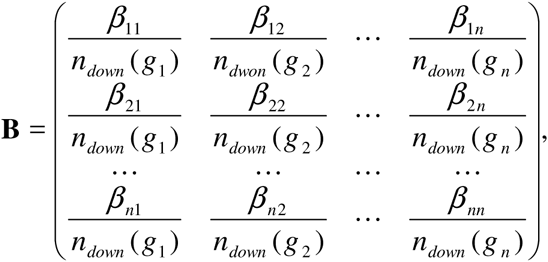

**I** is the identity matrix, and

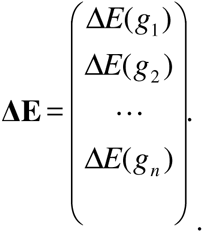

The overall score for pathway pertubation calculated as: 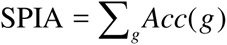.

### Statistical tests

Principal component analyses were performed using the MADE4 package [Culhane, 2005]. Hierarchical clustering heat maps with Pearson distances and average linkage were generated using heatmap.2 function from the *gplots* package [Scales, 2015].

### Mathematical modeling

We performed a Monte Carlo trial to investigate the data aggregation effect. Something may be missing here? We assumed that the number of genes in each pathway is distributed log-normally with the variable median number N. The case-to-normal-ratio (*CNR*) values for each gene were also sampled from the log-normal law, so that the value of log *CNR* had a normal distribution. When sampling *CNR*, we distinguished between *biased* and *unbiased* models of gene expression. For the *biased* model, the *CNR* distribution has a median value of 1.5, whereas for the *unbiased* model, the median *CNR* value is 1. The standard deviation of the mean log *CNR* value was set to 0.3 for both biased and unbiased models. The independent error produced by an experimental platform was also sampled stochastically. We simulated both the *exact* and *noisy* expression profiling methods. By the definition, *exact* methods did not introduce errors. For *noisy* methods, the error was chosen from the log-normal distribution, with a median value of 1.0. All the calculations were made using the R open source platform (version 3.1.2).

### Analysis of published transcriptomic and proteomic datasets

Prior to analysis, all the microarray data were quantile normalized [Bolstad, 2003], and the RNA-seq data were normalized using the DESeq package from Bioconductor software [Anders, 2010]. All gene products showing zero intensities were skipped to avoid aberrant data variations. Pearson correlation coefficients between the same samples examined using different expression profiling methods (e.g., proteome vs transcriptome or MH vs NGS) were calculated at two levels of data aggregation: first, at the level of distinct genes and gene products – namely for the value of log *CNR* (the so-called *C*_*g*_ correlation value); and, second, at the level of the whole pathways, for the *PAS* value (the ***C_p_*** correlation coefficient). Then, the ratio 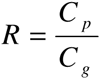 was calculated for each sample.

### Analysis of biological relevance after cross-platform harmonization

Transcriptional profiles were obtained using the Agilent Whole Human Genome Oligo and Affymetrix Human Genome U133 Plus 2.0 array platforms. The transcriptomic data were cross-platform harmonized with the XPN method [Shabalin, 2008] using the R package CONOR [Rudy, 2011]. Then, the cross-harmonized gene expression patterns between the Agilent and Affymetrix platforms were used as the input data for the *PAS* calculations. For all the calculations, type A samples were used as *normal*, and type B, C and D samples - as *cases*.

Euclidean distances between the *PAS* vectors were used to determine whether the resulting *PAS* samples are grouped in agreement with their biological properties (i.e., biological sample types B, C and D compared to A), or according to the experimental platform used to investigate them (i.e., Agilent or Affymetrix microarray platform). The cluster dendrograms and violin plots were drawn using the R packages *dendextend* and *vioplot*, respectively.

## ACKNOWLEDGEMENTS

This work was supported by the Pathway Pharmaceuticals Research Initiative (Hong Kong). The authors thank the First Oncology Research and Advisory Center (Moscow, Russia) for the support in preparation of this manuscript. We would like to thank Alex Kim and ASUS for equipment and support of this research.

## Footnotes

1 In the current paper, we apply the term *normalization* to any method for *within-platform* batch effect elimination, and *harmonization* when such procedure is performed for the *cross-platform* comparison, although the mathematical methods for both the former and latter procedures may be different.

